# Identifying Causal Relations between Gut Microbiome, Metabolic Dysfunction-Associated Fatty Liver Disease and the Novel Mediators of Blood Metabolites

**DOI:** 10.1101/2024.08.05.606637

**Authors:** Xinghao Yi, Haoxue Zhu, Mengyu He, Ling Zhong, Shan Gao, Ming Li

## Abstract

**Background:** Research has established connection between gut microbiome and the risk of metabolic dysfunction-associated fatty liver disease (MAFLD). However, the causal relationships and the roles of potential mediating factors, such as blood metabolites, remain unclear.

**Methods:** We conducted a bidirectional and mediation Mendelian randomization (MR) study using the genome-wide summary statistics of gut microbial taxa (Dutch Microbiome Project, n = 7,738), blood lipids (UK Biobank, n =8,299), and the largest MAFLD GWAS data (1,483 cases and 17,781 controls). We used the inverse-variance weighted estimation method as our primary approach. The multivariable Mendelian randomization (MVMR) and two-step MR approaches were used to prioritize the most likely causal metabolites as mediators. Additionally, we conducted linkage disequilibrium score regression (LDSC) analyses to assess genetic correlations, and downstream gene-based analyses to investigate the shared biological mechanism.

**Results:** By testing the causal effects of 205 bacterial pathways and 207 taxa on MAFLD, we identified 5 microbial taxa causally associated with MAFLD, notably the species Parabacteroides merdae (OR [95%CI] = 1.191[1.022-1.388], *p* = 0.025). Among 1,399 blood metabolites, 53 showed causal associations with MAFLD, with pregnenetriol sulfate identified as a mediator for genus Parabacteroides on MAFLD (proportion mediated = 16.30%). LDSC analysis also provided suggestive evidence for a potential genetic correlation between them (r_g_= 2.124, *p*=0.009).

**Conclusions:** The study suggested a novel causal relationship between gut microbial taxa and MAFLD, especially the genus Parabacteroides merdae and blood metabolite pregnenetriol sulfate might mediate this relationship.

**Importance:** Our study reveals novel insights into how the intersection of microorganisms living in the human gut, known as the gut microbiome, influences the development of Metabolic Dysfunction-Associated Fatty Liver Disease (MAFLD), a condition increasingly recognized as a major global health concern. By identifying specific gut microbiome taxa and metabolites that contribute to the onset and progression of MAFLD, our findings enhance comprehension of this prevalent condition and unveil promising prospects for its prevention and intervention. We discovered that certain gut bacteria can affect the levels of blood metabolites, which in turn impact the liver’s health. This work carries significant implications for novel strategies for MAFLD prevention and treatment, including interventions aimed at modifying the gut microbiome. Our research underscores the gut-liver connection and its implications for metabolic diseases, contributing to future therapeutic developments that could improve public health worldwide.

**Graphic abstract:** 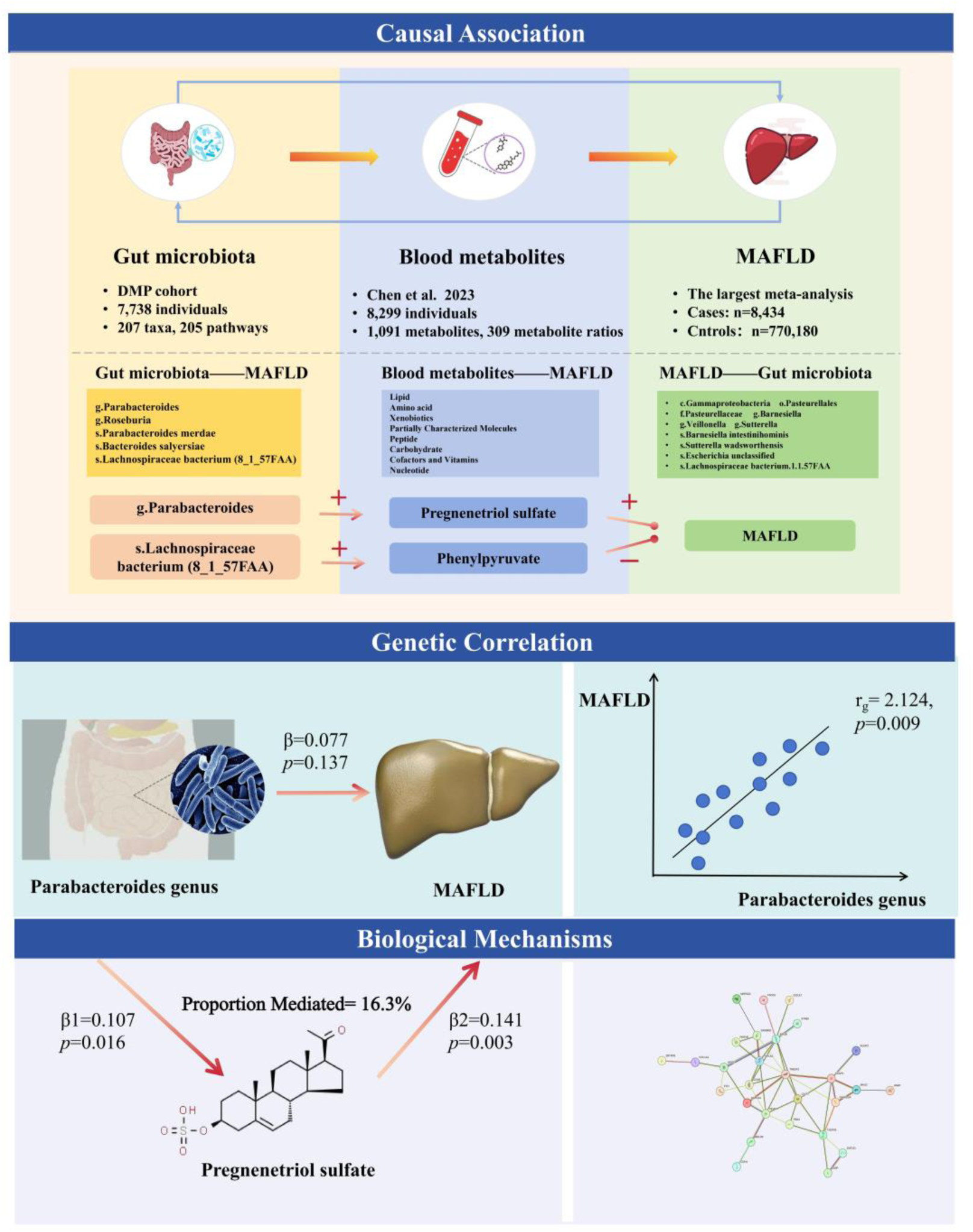

## 1. Introduction

Compelling evidence indicates that gut microbiota plays a significant role in several metabolism-related diseases, including obesity, type 2 diabetes, and fatty liver disease^1, 2^. Given that the liver predominantly derives its blood supply from the portal vein, it is particularly susceptible to influence originating in the gut. This includes the gut microbiota and the bioactive byproducts generated from the fermentation of food by these microbes. Furthermore, recent studies have highlighted the role of the gut microbiota and metabolites in the progression of fatty liver^3, 4^. Non-alcoholic fatty liver disease (NAFLD) ranks among the most common chronic liver conditions worldwide, impacting as many as 30% of people globally^5^. Often associated with obesity and insulin resistance, in 2020, it was renamed as metabolic dysfunction-associated fatty liver disease (MAFLD)^6^. To date, despite significant efforts to prevent or treat MAFLD, effective prevention strategies and treatments remain elusive.

Therefore, identifying modifiable risk factors is crucial to reduce the economic and social impact of MAFLD. Recent studies highlighting the connection between gut microbiota and metabolic-related diseases have positioned gut microbiota as a promising target for early prevention and clinical treatment. MAFLD has been associated with altered intestinal function^7^. For example, gut microbiota may alter intestinal permeability, leading to the release of lipopolysaccharides that can trigger systemic and tissue inflammation^8^. Several cohort studies indicate that patients with MAFLD exhibit unique microbial alterations in their composition and function^9, 10^. However, the evidence linking gut microbiota to MAFLD remains associative rather than causal.

Abnormal blood metabolites have been linked to an increased risk of MAFLD^11^. Plasma metabolomics has identified thousands of metabolites, many of which exhibit potential as biomarkers for precision medicine^12^. Recent epidemiological studies, animal experiments, and Mendelian randomization (MR) analysis consistently revealed correlations between the gut microbiota and the circulating metabolome^13, 14^. These correlations intimate that plasma metabolites could be instrumental in the etiology of MAFLD, potentially acting both as direct contributors to the disease process and as intermediaries within the intricate interplay between the gut microbiota and MAFLD. Nonetheless, definitive evidence elucidating the causal mechanisms linking the gut microbiota to MAFLD through alterations in blood metabolites remains to be established.

Previous robust genome-wide association studies (GWAS) have identified numerous single nucleotide polymorphisms (SNPs) associated with both MAFLD and gut microbiota, providing a unique chance to explore their potential causal connections by MR analysis. MR utilizes genetic variants (specifically SNPs) linked to specific risk factors as instrumental variables to infer causal relationships between these factors and health outcomes. This approach is similar to the randomized controlled trial methodology in clinical research, as it circumvents the biases introduced by reverse causality and confounding variables^15^.

In this research, we applied two-sample bidirectional and multivariable MR approaches to explore the causal links between gut microbiota and MAFLD, investigating blood metabolites as potential mediators. We further evaluated the genetic correlation by LDSC analysis. These approaches not only enhance our understanding of the complex interplay between the gut microbiota and MAFLD, but also pave the way for developing targeted therapeutic strategies that address the metabolic mediators involved.

## 2. Methods

### 2.1 Study design

The methodological framework of the current investigation is delineated in **Figure 1**. The research design has been systematically refined into three pivotal stages, which are elaborated upon as follows. Initially, we executed bidirectional univariable Mendelian randomization (UVMR) analyses to explore potential associations between the gut microbiome (207 taxa and 205 bacterial pathways) and MAFLD. Subsequently, we directed our focus toward 1,399 blood metabolites as candidate mediators in the association between gut microbiome and MAFLD using UVMR and MVMR analyses, with the mediating effects being quantified using a two-step MR approach. In the final stage, we conducted LDSC analyses to assess genetic correlations, and engaged in an extensive investigation encompassing Protein-Protein Interaction (PPI) analysis, pathway analysis, and Gene Ontology (GO) analysis to unravel the intricate mechanisms behind the identified mediation effects.

**Figure 1.**
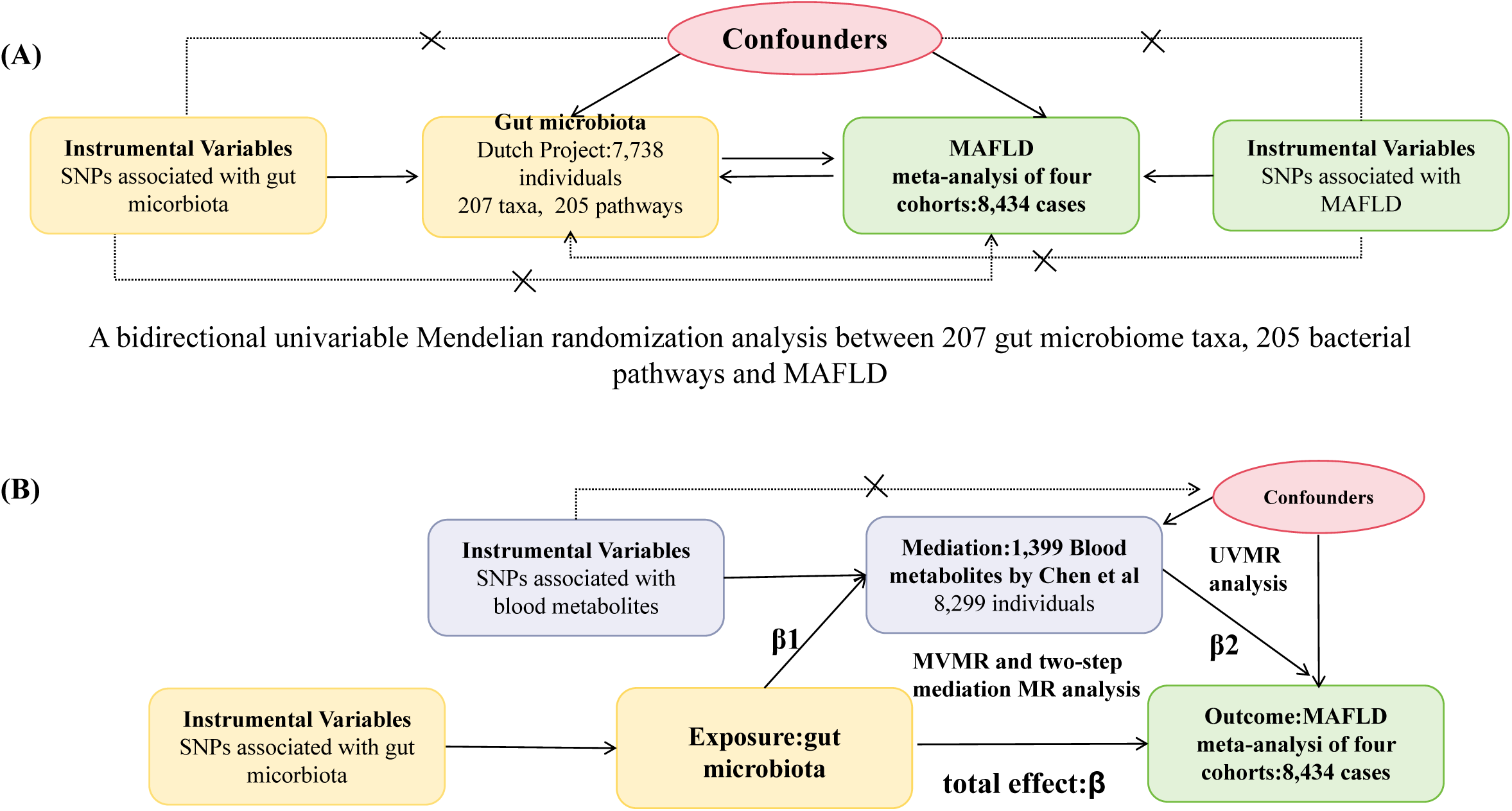
The overall study design. (A) A bidirectional univariable Mendelian randomization analysis between the 207 gut microbiome taxa and MAFLD, and 205 bacterial pathways and MAFLD; (B) A multivariable and two-step Mendelian randomization analysis of the effect of the gut microbiome on MAFLD via the blood metabolites. MAFLD, metabolic associated fatty liver disease.

### 2.2 Data source

The datasets utilized in the present study were exclusively sourced from publicly accessible repositories, and consequently, ethical approval was not deemed necessary.

The GWAS summary statistics pertaining to the gut microbiota were derived from the most recent and pioneering GWAS on the gut microbiome in a cohort of 7,738 participants from the Dutch Microbiome Project, including a total of 205 bacterial pathways and 207 taxa (5 phyla, 10 classes, 13 orders, 26 families, 48 genera, and 105 species)^16^.

The summary data on MAFLD were extracted from the largest meta-analysis, encompassing four European ancestry cohorts: the Electronic Medical Records and Genomics network, the UK Biobank, the Estonian Biobank, and the FinnGen consortium. This comprehensive analysis incorporated a total of 8,434 cases diagnosed with MAFLD and a robust control group comprising 770,180 individuals, as reported in a study^17^.

For the blood metabolites, we leveraged the GWAS summary statistics from a pivotal investigation led by Chen et al^18^. This study quantified 1,091 distinct metabolites and 309 metabolite ratios across a cohort of 8,299 participants, utilizing genome-wide genotyping to elucidate the genetic underpinnings of metabolic variations. The comprehensive summary of the data utilized in this study is listed in **Supplementary Table 1**.

### 2.3 Bidirectional UVMR analyses between the gut microbiome and MAFLD

We conducted a two-sample bidirectional UVMR to estimate the total effect of the gut microbiome on MAFLD. Three essential assumptions must be taken into account when selecting instrumental variables (IVs) for subsequent MR analyses^19^. These assumptions include the reliability and strength of the relationship between the IVs and the exposure, the absence of any confounding associations between the IVs and potential confounders, and the premise that the IVs exert their influence on the outcome exclusively through the exposure. In this research, we loosed the genome-wide significance threshold from 5×10^-8^ to 1×10^-5^ to identify additional IVs for gut microbiota and excluded SNPs in linkage disequilibrium (LD, R^2^ < 0.001) within a 10,000 kilobase pairs window.

In the UVMR analyses, the inverse–variance weighted (IVW) method was adopted as the principal approach for estimating causal effects, and four additional methods (MR Egger, Weighted median, Simple mode, and Weighted mode) were utilized to provide a comprehensive assessment. We calculated the odds ratio (OR) alongside its corresponding 95% confidence intervals (95%CI) to assess the impact of a 1-ln increment in the gut microbiota on the probability of developing MAFLD. A two-sided *p*-value that passed the Bonferroni correction was defined as statistically significant, which is 0.0002 (0.05/207) for microbial taxonomy and 0.0002 (0.05/205) for bacterial pathways, and a *p*-value < 0.05 but not meeting the Bonferroni-corrected threshold was considered to have a suggestively significant association.

### 2.4 Mediation MR Analyses

The methodologies of the selection of mediational variables are elaborately depicted in **Supplementary Figure 1**. We performed UVMR analyses between 1,399 blood metabolites and MAFLD and narrowed our focus to those metabolites that were causally associated with the disease, selecting them for further mediation analysis. MVMR and two-step UVMR analyses were both conducted to assess whether any intermediate metabolite could be mediating the relationship between microbial taxonomy and MAFLD^20^, and calculated mediation effects using the delta method^21^.

Two-step MR analysis was detailed as follows. The first step was performed to estimate the causal effect of gut microbial taxonomy on the blood metabolites mediator (β1) using UVMR and the effect should be unidirectional. The second step was to estimate the causal effect of the mediator on MAFLD with adjustment for gut microbial taxonomy (β2) using MVMR. Subsequently, the mediated proportion of the total effect of gut microbial taxonomy on MAFLD through each mediator was determined by calculating the indirect effect, derived from the above two steps (indirect effect: β1×β2), and dividing it by the total effect.

The UVMR analysis between 1,399 blood metabolites and MAFLD selected a two-sided *p*-value less than 3.57ⅹ10^-5^ (0.05/1,399, Bonferroni-corrected significance) as statistically significant. Nevertheless, in pursuit of more comprehensive mediators, we also considered a less conservative threshold of 0.05 as an alternative criterion. The first-step UVMR estimates of microbial taxonomy on metabolites were represented by β with 95%CI.

### 2.5 MR Sensitivity analyses

We conducted the Cochran’s Q statistic to quantify the heterogeneity effects among the selected SNPs, and the MR-Egger regression test to assess the pleiotropic effects. The *p*-values exceeding 0.05 indicate the absence of heterogeneity and horizontal pleiotropy. Additionally, leave-one-out analysis was performed to evaluate the individual influence of a single SNP on the outcome, by removing each SNP one by one and reassessing the causal effects. All statistics were calculated using the R package “TwoSampleMR” (version 0.5.7) in R software (version 4.2.2).

### 2.6 Genetic correlation analysis

We utilized LDSC to estimate the genetic correlation (rg) between MAFLD and gut microbiome, which have been previously identified to have casual associations with MAFLD through MR analysis. This methodological approach involves an examination of the relationship between test statistics and linkage disequilibrium, which serves to quantify the extent of inflation attributable to either a genuine polygenic signal or potential bias^22^. Specifically, the process involves multiplying the z-scores of each variant associated with Trait 1 by the corresponding z-scores of each variant from Trait 2. Subsequently, the genetic covariance is derived from a regression analysis of this product against the LD score. The genetic correlation is then ascertained by normalizing the genetic covariance by the heritability attributable to SNPs.

In this study, statistical significance was defined by a *p-value* of less than 0.01, following Bonferroni correction for multiple comparisons (0.05/5). A *p-value* ranging from 0.01 to 0.05 was interpreted as providing suggestive genetic correlation.

### 2.7 Biological annotation

Following the elucidation of the “gut microbial taxonomy-blood metabolites-MAFLD” pathways, we advanced our investigation by identifying the corresponding genes associated with these traits from GWAS summary statistics. We then delved into the intricacies of these pathways by identifying PPI networks constructed using the mediation pathways and employing STRING database version 11.5^23^. We focused on the multiple interactive genes and conducted pathway and GO analysis.

## 3. Results

### 3.1 Bidirectional UVMR analysis of microbial taxa, bacterial pathways and MAFLD

The detailed characteristics of SNPs associated with gut microbiome (taxa and bacterial pathways), blood metabolites, and MAFLD can be found in **Supplementary Tables 2–4**, respectively.

In the forward UVMR analysis, a total of 207 taxa and 205 bacterial pathways were assessed for their associations with MAFLD. Based on the IVW method, five microbial taxa demonstrated significant correlations, with *p*-values below the threshold of 0.05. Among these, four taxa exhibited a positive association with the risk of developing MAFLD (**Figure 2A**), including the genera Parabacteroides (OR[95%CI] = 1.122[1-1.258], *p* = 0.05) and Roseburia (OR[95%CI] = 1.147[1.03-1.276], *p* = 0.012), as well as the species Parabacteroides merdae (OR[95%CI] = 1.191[1.022-1.388], *p* = 0.025) and Bacteroides salyersiae (OR[95%CI] = 1.082[1.011-1.158], *p* = 0.023). Besides, one Lachnospiraceae bacterium (8_1_57FAA) at the species level presented a negative correlation with the risk of MAFLD (OR [95%CI] = 0.946[0.897-0.998], *p* = 0.042). The results by additional MR methods including MR Egger, weighted median, weighted mode, and simple mode are presented in **Supplementary Table 5**.

**Figure 2.**
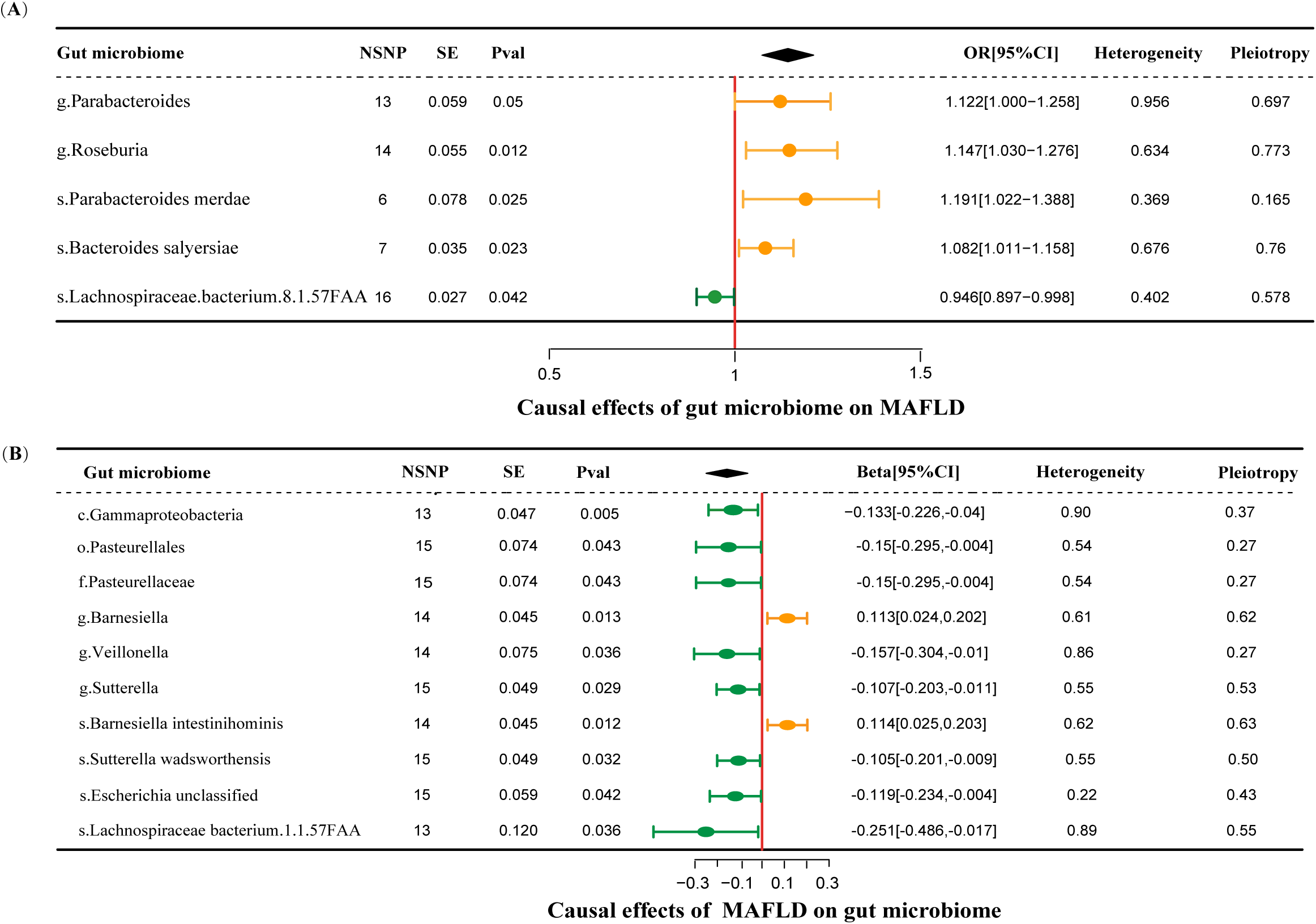
Forestplot of bidirectional causal effects between gut microbial taxa and MAFLD (*p* < 0.05). (A) Causal effects of gut microbial taxa on MAFLD; (B) Causal effects of MAFLD on gut microbial taxa. NSNP, number of SNPs; SE, standard error of coefficient estimate; Pval, the *p-value* of causal estimation in the IVW method; OR [95%CI], odds ratio and 95% confidence interval; Heterogeneity, the *p-value* of heterogeneity analysis; Pleiotropy, the *p-value* of horizontal pleiotropy analysis.

In the reverse MR analysis encompassing 205 gut microbial taxa in relation to MAFLD, we identified 10 significant casual associations (**Figure 2B**). The condition MAFLD demonstrated a negative correlation with eight taxa, while it was positively correlated with two others. Specifically, at the taxonomic levels of class, order, and family, MAFLD was associated with a diminished abundance of Gammaproteobacteria, Pasteurellales, and Pasteurellaceae, respectively. When examining the genus level, MAFLD was found to reduce the prevalence of Veillonella and Sutterella, conversely, it was linked to an increased abundance of Barnesiella. Furthermore, at the species level, there was a decrease in the abundance of Sutterella wadsworthensis, the unclassified Escherichia, and Lachnospiraceae bacterium (1_1_57FAA), whereas the abundance of Barnesiella intestinihominis was elevated. **Supplementary Figure 2** illustrates the MR scatter plots for both the forward and reversed MR analyses. Sensitivity analyses, including the Q statistics from the inverse-variance weighted (IVW) test and the MR-Egger regression, revealed no significant heterogeneity or horizontal pleiotropy, thus substantiating the robustness of our findings. The results by additional MR methods are presented in **Supplementary Table 6**.

In the realm of microbiota metabolic pathways, our analysis revealed that an increase in the abundance of ten specific pathways, quantified per a single unit increment, is causally associated with MAFLD, including six protective pathways (ORs < 0.01) and four risk pathways (ORs > 0.01) (**Supplementary Figure 3A**). In the reverse MR analysis, MAFLD was casually associated with seven pathways (**Supplementary Figure 3B**). These causal associations have withstood rigorous sensitivity analyses, demonstrating their robustness and reliability.

### 3.2 Blood metabolites as candidate mediators between gut microbial taxa and MAFLD

#### 3.2.1 Causal associations of blood metabolites with MAFLD

We conducted UVMR analyses between a panel of 1,399 blood metabolites and MAFLD and pinpointed 53 metabolites that exhibited a causal association with MAFLD (**Figure 3, Supplementary Table 7**). This diverse set of metabolites encompassed 19 distinct lipid species, 13 amino acids, 3 xenobiotics, a duo of carbohydrates and nucleotides, as well as partially characterized molecules and metabolite ratios. Additionally, our findings included a single cofactor and vitamin, a unique peptide, and eight metabolites whose identities remain to be elucidated. This result was supported by the MR sensitivity analyses (**Supplementary Table 8**).

**Figure 3.**
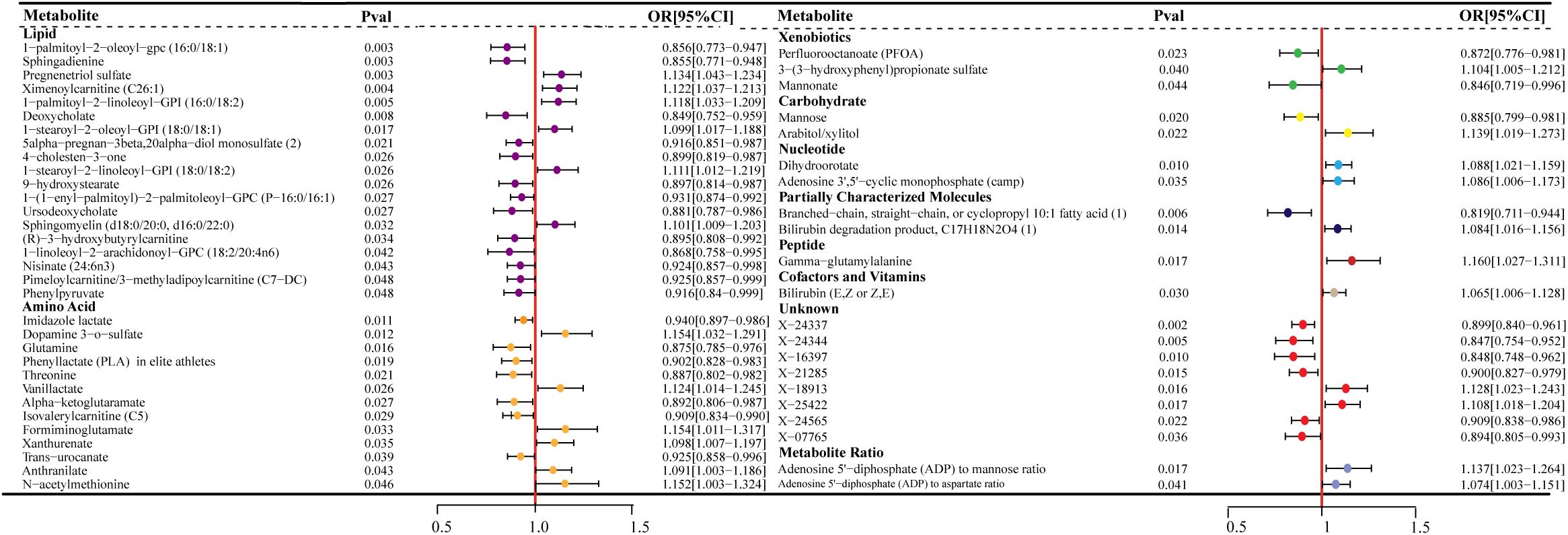
Forestplot of causal effects of blood metabolites on MAFLD (*p* < 0.05). Plasma metabolites encompass a diverse array of substances, including lipids, amino acids, xenobiotics, nucleotides, cofactors, and vitamins, as well as carbohydrates, peptides, partially characterized molecules, and unidentified compounds. Additionally, two metabolite ratios are identified. Pval, the *p-value* of causal estimation in the IVW method; OR[95%CI], odds ratio and 95% confidence interval.

#### 3.2.2 Two-step mediation MR analysis between 5 gut microbial taxa, 53 blood metabolites and MAFLD

In the first step of estimating causal effects between the above 5 MAFLD-related microbial taxa and 53 MAFLD-related metabolites using UVMR, 11 out of 53 candidate metabolite mediators were shortlisted (**Table 1**). The Parabacteroides genus was found to positively influence the levels of pregnenetriol sulfate and X-07765, while the Roseburia genus exhibited a negative effect on xanthurenate concentrations. Specifically, the species Parabacteroides merdae was associated with increased levels of branched-chain, straight-chain, and cyclopropyl 10:1 fatty acid, as well as X-07765. Bacteroides salyersiae was linked to elevated levels of 3-(3-hydroxyphenyl) propionate sulfate, anthranilate, and X-18913, but showed a decrease in X-07765 levels. Lachnospiraceae.bacterium.8.1.57FAA was observed to enhance phenylpyruvate levels and reduce alpha-ketoglutaramate levels.

**Table 1.** The MR analysis of gut microbial taxa on the blood metabolites.

| Exposure: gut microbiome | Outcome: blood metabolites | nSNP | Se | p-value | Beta [95%CI] |
| --- | --- | --- | --- | --- | --- |
| g.Parabacteroides | Pregnenetriol sulfate | 14 | 0.045 | 0.016 | 0.107[0.019-0.195] |
| g.Parabacteroides | X-07765 | 14 | 0.053 | 0.010 | 0.137[0.033-0.241] |
| g.Roseburia | Xanthurenate | 14 | 0.055 | 0.029 | -0.119[-0.227--0.011] |
| s.Parabacteroides merdae | Branched-chain, straight-chain,or cyclopropyl 10:1 fatty acid (1) | 7 | 0.064 | 0.037 | 0.134[0.009-0.259] |
| s.Parabacteroides merdae | X-07765 | 7 | 0.081 | 0.026 | 0.181[0.022-0.34] |
| s.Bacteroides salyersiae | 3-(3-hydroxyphenyl)propionate sulfate | 8 | 0.037 | 0.012 | 0.094[0.021-0.167] |
| s.Bacteroides salyersiae | Anthranilate | 8 | 0.037 | 0.039 | 0.076[0.003-0.149] |
| s.Bacteroides salyersiae | X-07765 | 8 | 0.031 | 0.030 | -0.068[-0.129--0.007] |
| s.Bacteroides salyersiae | X-18913 | 8 | 0.031 | 0.018 | 0.073[0.012-0.134] |
| s.Lachnospiraceae.bacterium.8.1.57FAA | Alpha-ketoglutaramate | 16 | 0.024 | 0.023 | -0.055[-0.102--0.008] |
| s.Lachnospiraceae.bacterium.8.1.57FAA | Phenylpyruvate | 16 | 0.025 | 0.032 | 0.053[0.004-0.102] |

**Abbreviations:** nSNP, number of SNPs; Se, standard error of coefficient estimate; *p*-value, *p*-value of causal estimation in the IVW method; Beta [95%CI], estimated causal effect coefficient and 95% confidence interval.

Having identified 11 potential mediators, we performed MVMR analyses to ascertain whether the causal association between each mediator and MAFLD remained consistent with adjustment for the selected gut microbial taxa. Four causal relationships between several metabolites and MAFLD persisted, including pregnenetriol sulfate, xanthurenate, branched-chain, straight-chain, or cyclopropyl 10:1 fatty acid (1) levels, and phenylpyruvate. We presented the comprehensive findings of the MVMR analysis in **Supplementary Table 9**. Subsequently, we refined our selection of mediators by assessing the consistency of the observed effect directions. The congruence between the total effect of gut microbial taxa on MAFLD and their indirect effects through specific metabolites (β1ⅹβ2, where β1 represents the causal effect of gut microbial taxa on metabolites, and β2 represents the causal effect of metabolites on MAFLD) allowed us to narrow down the potential mediators. Ultimately, we identified two metabolites that met all established criteria, thereby designating them as the final mediators. The reverse MR analysis validation between these 2 metabolites, gut microbiota, and MALFD was performed to ensure that the causal effects are unidirectional, the details are listed in **Supplementary Table 10**. Notably, the Parabacteroides genus exerted detrimental effects on MAFLD by increasing pregnenetriol sulfate levels, with a mediated proportion of 16.3%, while the Lachnospiraceae.bacterium.8.1.57FAA species exerted a protective effect on MAFLD, mediated by its enhancement of phenylpyruvate levels, which are shown to reduce the risk of developing the disease, accounting for a mediated proportion of 9.68% (**Figure 4**).

**Figure 4.**
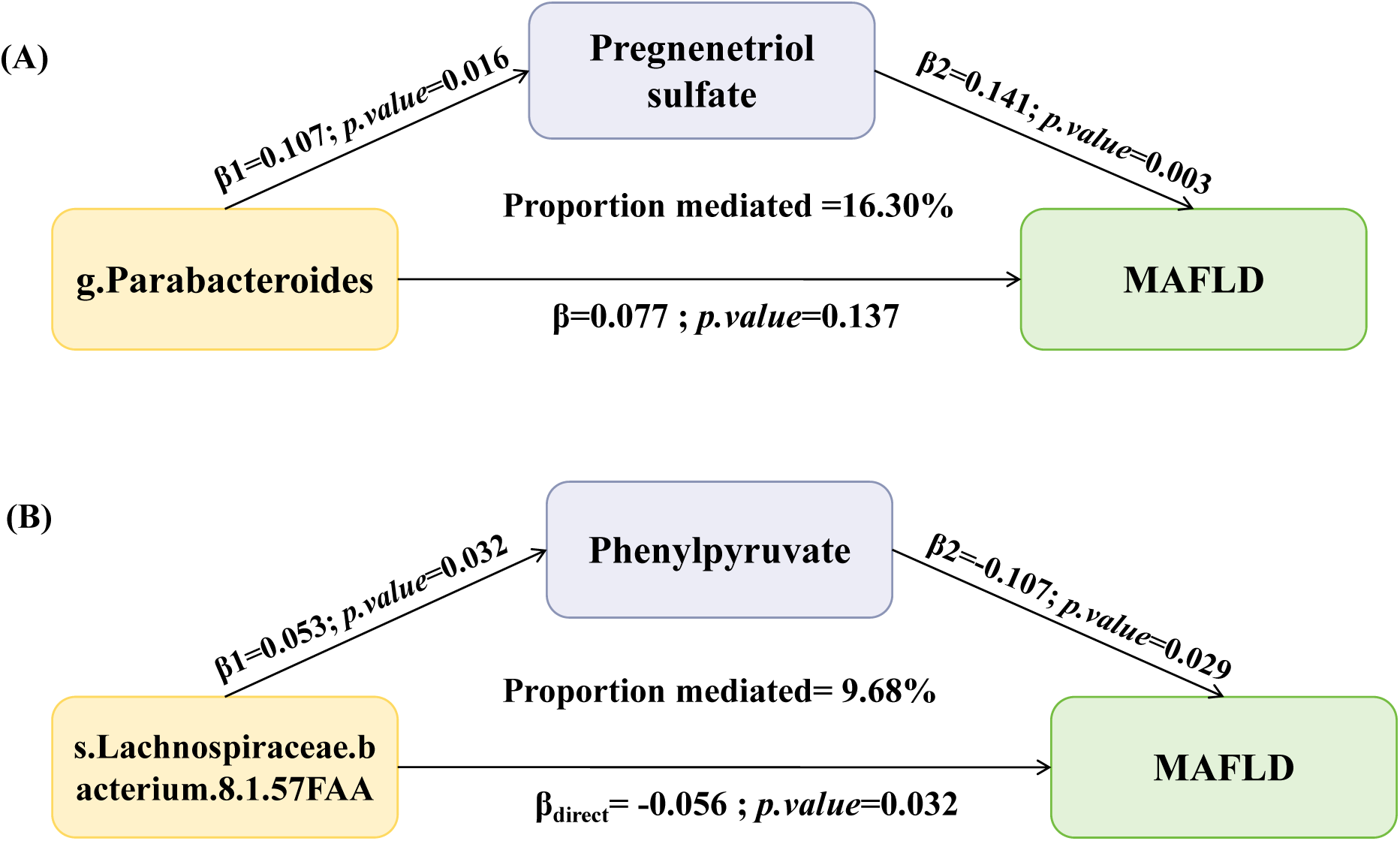
The mediation effect of gut microbial taxonomy on MAFLD via blood metabolite. (A) The pregnenetriol sulfate mediated the causal effect of the Parabacteroides genus on MAFLD; (B) The phenylpyruvate mediated the causal effect of the Lachnospiraceae.bacterium.8.1.57FAA species on MAFLD. β1, the effect of gut microbial taxonomy on the blood metabolites; β2, the effect of the blood metabolite on MAFLD after adjustment for gut microbial taxonomy; β_direct_, the direct effect of the gut microbial taxonomy on MAFLD; β_total_, the total effect of the gut microbial taxonomy on MAFLD; *p.value*, the *p-value* of causal estimation; Proportion mediated (%), the proportion of effect mediated through blood metabolites and 95% confidence interval.

### 3.2 LDSC regression analysis

We conducted LDSC analysis to assess genetic correlations between 5 gut microbiota taxa and MAFLD. The Parabacteroides genus was found significant genetic correlation with MAFLD (r_g_= 2.124, *p*=0.009), while the Roseburia genus (*p*=0.382) and Bacteroides salyersiae species (*p*=0.249) demonstrated no potential genetic correlation. Owing to limitations such as low heritability and sample size, the Lachnospiraceae.bacterium.8.1.57FAA and Parabacteroides merdae species cannot be used for the above analysis.

### 3.3 The PPI network, pathway, and GO analysis explored potential biological mechanisms

The PPI network analysis was conducted for the aforementioned two “gut microbial taxa-blood metabolites-MAFLD” pathways. For the “Parabacteroides genus-Pregnenetriol sulfate levels-MAFLD” pathway (**Figure 5A**), we meticulously selected and included a total of 8, 25, and 52 genes for in-depth analysis. Remarkably, this network revealed 29 genes that exhibited correlations with other genes with enrichment *p-values* significantly below < 1.0ⅹ10^-16^. Among them, five genes including *DOCK7*, *APOB*, *FADS2*, *STAB2*, and *MRPS22* belong to pregnenetriol sulfate. These genes were found to predominantly interact with MAFLD through the gene *APOB*.

**Figure 5.**
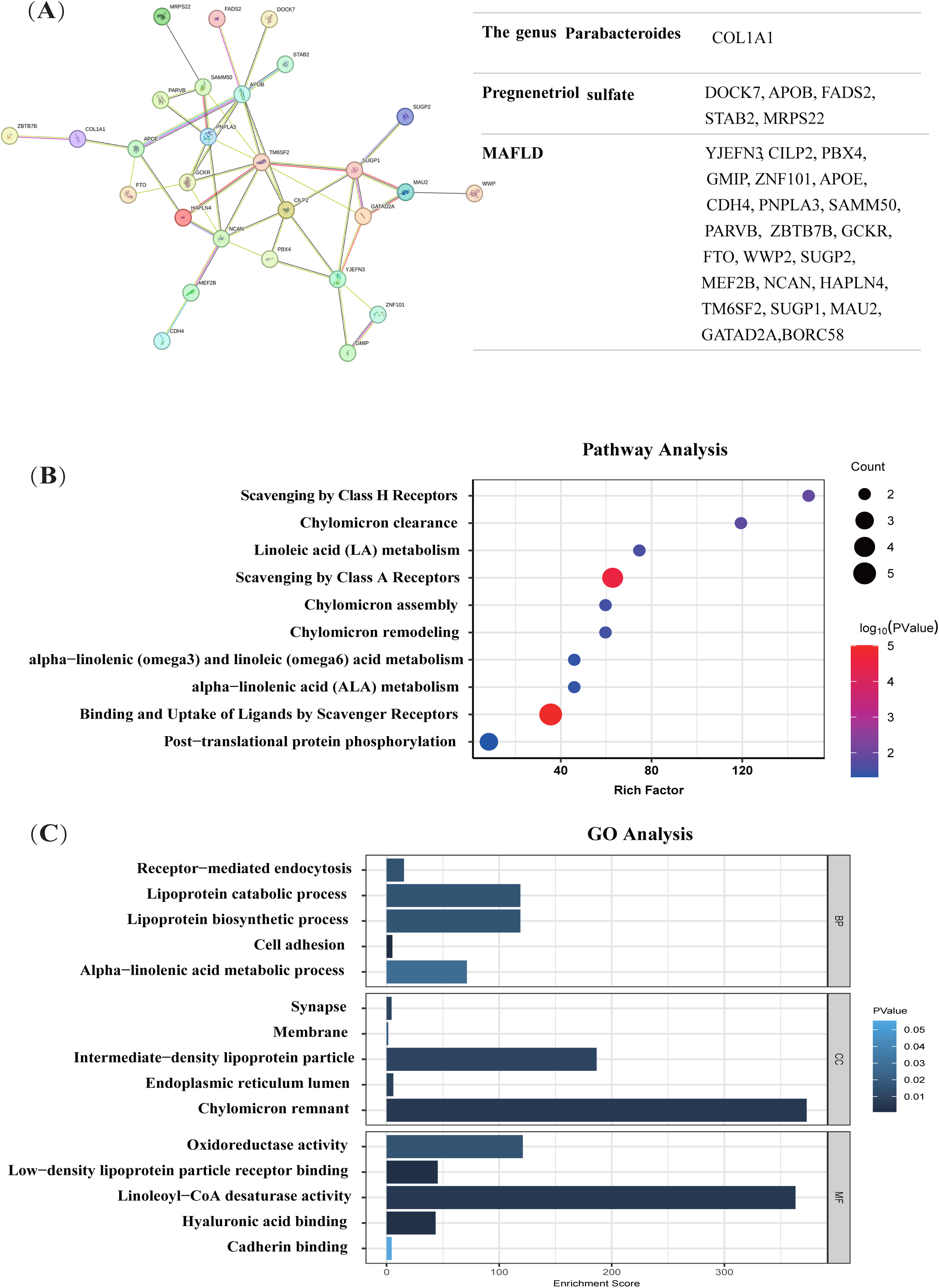
The “Parabacteroides genus-Pregnenetriol sulfate levels-MAFLD” pathway biological annotation. (A) The PPI network analysis of the Parabacteroides genus,pregnenetriol sulfate levels, and MAFLD. (B) The pathway analysis of multiple interactive genes in the “g.Parabacteroide -Pregnenetriol sulfate-MAFLD” pathway. (C) The GO analysis of multiple interactive genes in the “g.Parabacteroides-Pregnenetriol sulfate-MAFLD” pathway.

Only one gene, *COL1A1*, was retained and came from the genus Parabacteroides. The other remaining genes were involved in the MAFLD-associated genes. In the subsequent pathway and GO analysis, our findings revealed that the genes under investigation partake in a multitude of lipid-related biological processes. These encompass the assembly, remodeling, and clearance of chylomicrons, as well as the metabolism of linoleic acid and alpha-linolenic acid, which includes omega-3 and omega-6 fatty acid metabolism. Additionally, these genes were found to be associated with the metabolism of lipoproteins, specifically low-density lipoproteins and intermediate-density lipoproteins. The enzymatic activity of linoleoyl-CoA desaturase was also identified as a significant functional aspect of these genes (**Figures 5B and C**).

For the “s. Lachnospiraceae.bacterium.8.1.57FAA-Phenylpyruvate levels-MAFLD” pathway, 9, 21, and 52 genes were meticulously preserved for biological annotation analysis. The process ultimately led to the identification of 21 genes. However, a singular gene, *NUDT3*, which is associated with s. Lachnospiraceae.bacterium.8.1.57FAA and connected to MAFLD-related genes through *FTO*. Additionally, no genes related to phenylpyruvate metabolism were retained (**Supplementary Figure 4**). Furthermore, our pathway and GO analyses did not reveal any biological pathways significantly associated with this ensemble of genes.

## 4. Discussion

In the present large-scale MR study, 5 gut microbiota taxa and 53 blood metabolites were found to be causally associated with MAFLD. Regarding a possible underlying mechanism, among the 53 MAFLD-related blood metabolites, we uncovered 11 blood metabolites associated with the 5 microbiota taxa and MAFLD using MVMR as mediation analyses. Two significant mediating roles of blood metabolites were highlighted. Specifically, metabolites pregnenetriol sulfate and phenylpyruvate were identified as mediators, linking microbial taxa like Parabacteroides and Lachnospiraceae bacterium 8_1_57FAA to MAFLD. These findings suggest a pathway where microbial dysbiosis leads to altered metabolic profiles, which in turn contribute to liver disease progression.

The forward MR analysis identified significant associations between five specific microbial taxa and MAFLD. Notably, genera such as Parabacteroides and Roseburia, along with the species Parabacteroides merdae and Bacteroides salyersiae, were positively correlated with an increased risk of MAFLD. These findings align with studies suggesting that certain gut microbes may influence liver health through mechanisms like modulating bile acid metabolism and systemic inflammation^24^. Conversely, a Lachnospiraceae bacterium demonstrated a protective role, potentially by producing short-chain fatty acids (SCFAs) that improve insulin sensitivity and thus reduce hepatic steatosis^25, 26^. In the reverse MR analysis, relationships between MAFLD and ten microbial taxa were elucidated. This included a negative correlation with the abundance of taxa such as Gammaproteobacteria, Pasteurellales, and Veillonella, previously reported to be associated with liver diseases, especially carcinoma and cirrhosis^27, 28^. These associations suggest a potential protective role of these microbes against MAFLD, possibly through pathways involving the modulation of inflammation or the metabolism of harmful substances within the gut^29, 30^. Conversely, an increase in taxa like Barnesiella was linked to a higher risk of developing MAFLD, which could be attributed to their role in unfavorable metabolic processes impacting liver health^31^. Specifically, 4 species, namely Barnesiella intestinihominis, Sutterella wadsworthensis, Escherichia (unclassified), and Lachnospiraceae bacterium.1.1.57FAA were discovered, these insights underscore the complex interplay between the gut microbiome and MAFLD, pointing towards potential therapeutic targets for managing this condition. For instance, manipulating the gut microbiota through diet, probiotics, or antibiotics might offer novel approaches to prevent or treat MAFLD^30, 32^.

We identified pregnenetriol sulfate as a key mediator in the causal link between the genus Parabacteroides and MAFLD. Research has highlighted the crucial role of Parabacteroides in the progression of fatty liver, especially species like Parabacteroides distasonis, which can utilize dietary inulin to suppress non-alcoholic steatohepatitis (NASH) through its metabolite, pentadecanoic acid^33, 34^. However, these effects might be strain-specific and not necessarily representative of the entire genus, which is known to have dual potential for pathogenicity and probiotic effects^35^. Our study reveals that the genus Parabacteroides is positively associated with MAFLD risk, suggesting that the interactions of Parabacteroides with other gut microbiota could alter its role under certain conditions. Further research into the specific strains present and their metabolic functions in different host environments is essential to fully elucidate these complex interactions^24^. PPI analysis revealed that the genus Parabacteroides may function through MAFLD-related genes via *COLA1*, which encodes type I collagen alpha 1, a key component of the liver’s extracellular matrix. This gene is involved in epithelial-mesenchymal transformation, closely associated with the development of malignant tumors^36, 37^. Its role in MAFLD pathophysiology could be critical, particularly through its involvement in the structural and functional integrity of the liver. Type I collagen is vital for maintaining liver architecture and function^38^. In the context of MAFLD, the dysregulation of *COLA1* expression can lead to increased collagen deposition, contributing to liver fibrosis, which is a significant and potentially irreversible step in the progression of MAFLD to more severe stages such as cirrhosis and hepatocellular carcinoma^39^. Understanding how pregnenetriol sulfate influences *COLA1* expression could provide insights into the hormonal and metabolic pathways that are dysregulated in MAFLD and offer new avenues for therapeutic interventions aimed at modulating these pathways to prevent or mitigate fibrosis. Pregnenetriol sulfate, a steroid metabolite and specifically a sulfate ester of pregnenetriol, is involved in various biological processes, including hormonal activity and metabolism^40, 41^. A study discovered that lower pregnenetriol sulfate levels were observed in plasma from women who subsequently developed breast cancer^42^. The interaction of pregnenetriol sulfate with *COLA1* (represents of genus Parabacteroides) suggested a possible mechanism where hormonal or metabolic imbalances induced by changes in steroid metabolism could influence the expression and activity of type I collagen. This interaction may exacerbate the fibrotic process by promoting the deposition of collagen fibers, thus enhancing the fibrogenic response within the liver. To sum up, the exact functions and mechanisms of pregnenetriol sulfate in the body, especially in relation to metabolic conditions like MAFLD, are areas of ongoing research. Its role could be further compared with other known influences on MAFLD, like changes in bile acids or short-chain fatty acid profiles linked to different gut bacteria^43, 44^. Such comparisons might clarify the unique or synergistic effects of pregnenetriol sulfate in the context of gut-liver axis dynamics.

Specious Lachnospiraceae.bacterium.8.1.57FAA was previously reported to be associated with estrogen, its’ abundance increased in mice with ovariectomy^45^. This is the first to report its association with MAFLD. Phenylpyruvate is primarily known as an intermediate in the catabolic pathway of phenylalanine^46^. This pathway is crucial for the breakdown and utilization of phenylalanine for energy production in the liver. This metabolite is linked to atherosclerotic cardiovascular disease and increased phenylpyruvate may augment inflammatory responses^47, 48^. Our findings reveal a paradoxical protective role of phenylpyruvate against MAFLD mediated by Specious Lachnospiraceae bacterium 8.1.57FAA. This suggests that the bacterium’s specific metabolic activities could modulate phenylalanine metabolism, potentially reducing the inflammatory and atherogenic effects of phenylpyruvate. The observed increase in this bacterium post-ovariectomy hints at a complex interaction between hormonal changes and gut microbiota composition, suggesting that estrogen may regulate the activity of this bacterium and influence the metabolic outcomes of phenylpyruvate. The enzymatic pathways facilitated by Specious Lachnospiraceae bacterium 8.1.57FAA that lead to phenylpyruvate production and their regulation under varying physiological states may need further exploration.

Key strengths of our study include the extensive genetic summary data utilized for blood metabolites and MAFLD, detailed species-level gut microbiota analysis, and the use of MVMR. This methodology enabled the effective prioritization of metabolites that show the strongest associations, enhancing the robustness of our findings. Subsequent LDSC analysis of the correlation between gut bacteria and MAFLD provided genetic substantiation for our MR discoveries, further fortifying the validity of our results with genetic evidence. However, we must acknowledge certain limitations of our study. Firstly, while MR offers valuable insights by simulating lifelong genetic influences, it may not fully capture the effects of short-term microbiome changes. Nonetheless, MR provides preliminary data on the directionality of these effects, setting the stage for more detailed investigations through animal studies and clinical trials. Secondly, our study’s sample size, though the largest to date for gut microbiome GWAS with species-level analysis, might still be insufficient for detecting all causal links.

Our research indicates that focusing on the gut microbiota genus Parabacteroides may offer a viable approach for MAFLD prevention in a clinical setting. Strategies could include the use of treatments like small-molecule antibiotics, engineered commensal or probiotic bacteria, prebiotics, and bioactive metabolites to adjust its abundance^32, 49^.

## 5. Conclusion

The present study, leveraging a comprehensive MR approach, has delineated a significant causal association between specific gut microbial taxa and MAFLD, with blood metabolites identified as pivotal mediators in this association. Our findings underscore the role of the gut microbiome in modulating metabolic pathways that influence the pathogenesis of MAFLD, providing a robust framework for future research and therapeutic interventions.

The identification of metabolites such as pregnenetriol sulfate and phenylpyruvate as mediators offers novel insights into the gut-liver axis and its complex interplay in the context of MAFLD. These discoveries prompt further investigation into the functional roles of these metabolites and their potential as targets for therapeutic modulation.

While our study’s extensive use of UVMR, MVMR, and two-step MR methodologies strengthen the causal inferences drawn, we acknowledge the necessity for continued research to fully elucidate the dynamic interactions within the gut microbiome and their metabolic consequences. The prospect of modulating gut microbial composition through dietary changes, probiotics, or targeted pharmacological interventions presents an exciting frontier for MAFLD prevention and management.

## 8. Authorship Contribution Statement

Xinghao Yi and Haoxue Zhu: Data curation, Formal analysis, Investigation, Software, Validation, Visualization, Writing original draft. These authors contribute equally to the article. Mengyu He and Ling Zhong: Data collection, Validation. Ming Li and Shan Gao: Funding acquisition, Investigation, Methodology, Project administration, Supervision, Writing – review & editing.

## 9. Data Availability

The GWAS summary statistics for gut microbiota can be downloaded in the GWAS Catalog under accession codes from GCST90027801 to GCST90027478 (https://www.ebi.ac.uk/gwas/efotraits/). The GWAS summary statistics for MAFLD were extracted from the largest meta-analysis with code GCST90091033 (https://www.ebi.ac.uk/gwas/studies/GCST90091033). The GWAS summary statistics for 1,399 blood metabolites were downloaded under accession codes from GCST90200429 to GCST90200288 (https://www.ebi.ac.uk/gwas/publications/36635386).

Data code will be made available on request.

## 10. Supplementary Materials

**Supplementary Table 1.** Detailed information for genome-wide association studies involved in the present Mendelian randomization study.

**Supplementary Table 2.** The instruments of gut microbial taxa. **Supplementary Table 3.** The instruments of gut bacterial pathways. **Supplementary Table 4.** The instruments of MAFLD.

**Supplementary Table 5.** The additional MR methods on the causal effect between 5 gut microbiome taxa and MAFLD.

**Supplementary Table 6**. The reverse MR analysis results between MAFLD and gut microbiome taxa.

**Supplementary Table 7.** The IVW MR analysis on the causal effect between 53 blood metabolites and MAFLD.

**Supplementary Table 8.** The sensitivity analysis results of MR analysis of 53 blood metabolites on MAFLD.

**Supplementary Table 9.** The MVMR analysis results of gut microbiota, blood metabolites and MAFLD.

**Supplementary Table 10.** The reverse MR analysis between two mediation pathways.

**Supplementary Figure 1.**
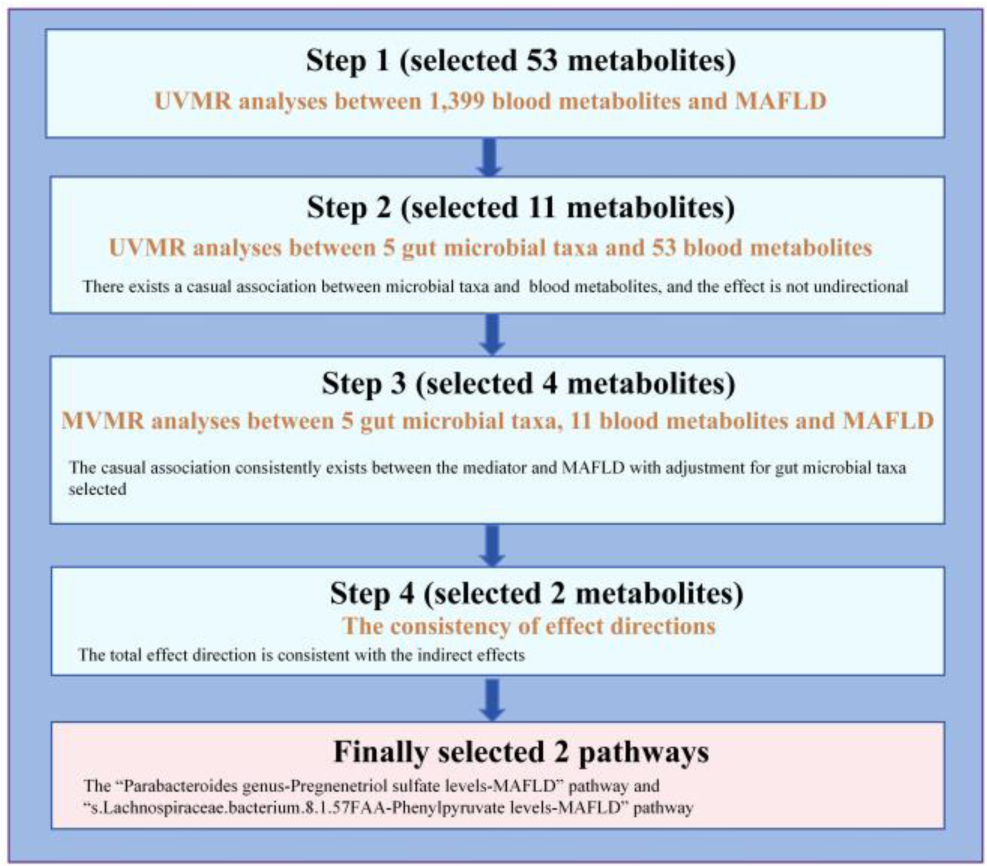
Mediator selection process in stage 2. We first screened candidate mediators for the association between gut microbial taxonomy and MAFLD by stringent criteria, and then calculated their mediating effects using two-step MR.

**Supplementary Figure 2.**
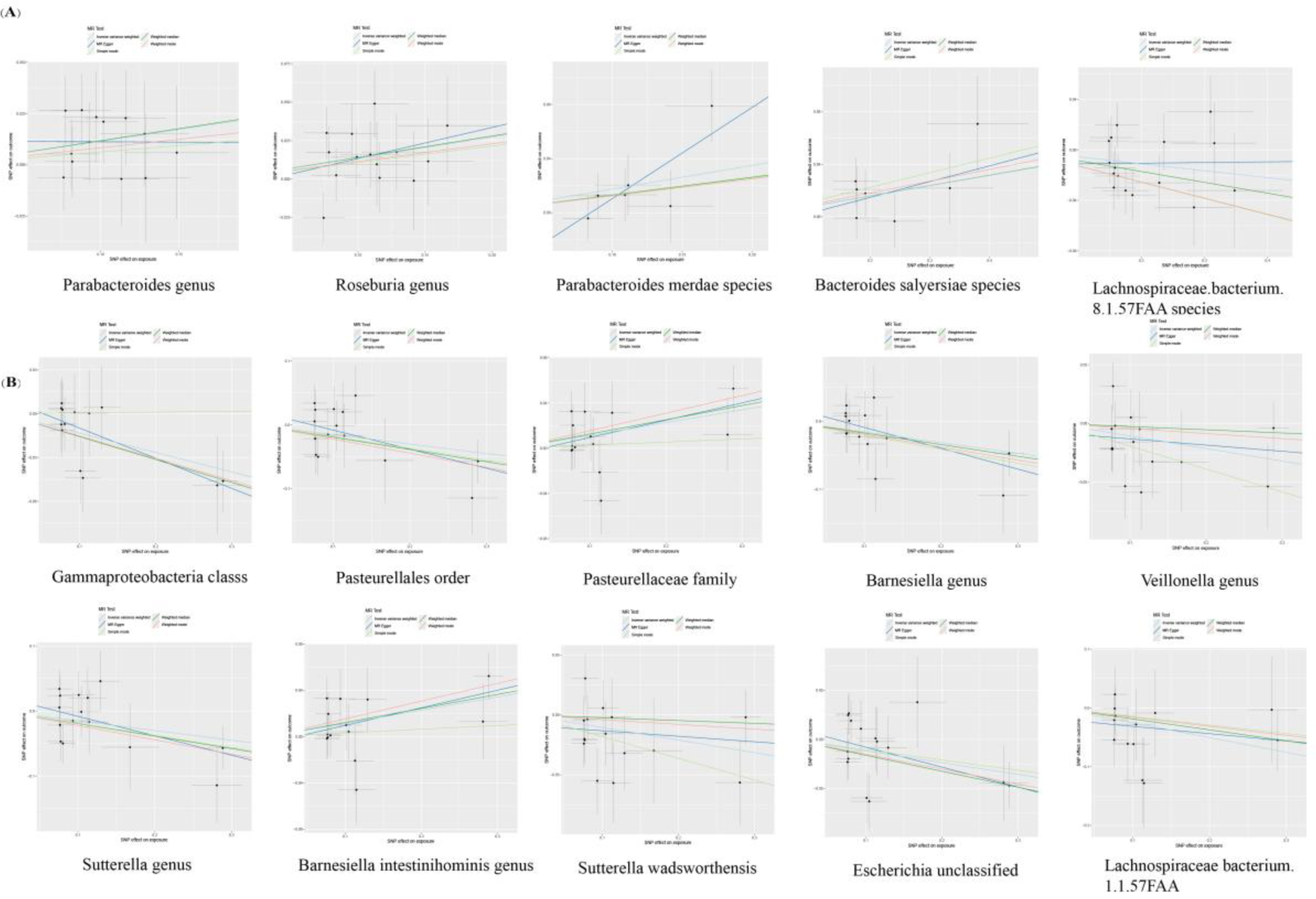
MR scatter plots. (A) Scatterplot of SNP potential effects of gut microbiome taxa on MAFLD; (B) Scatterplot of SNP potential effects of MAFLD on gut microbiome taxa. The slope of each line corresponds to the estimated MR effect per method.

**Supplementary Figure 3.**
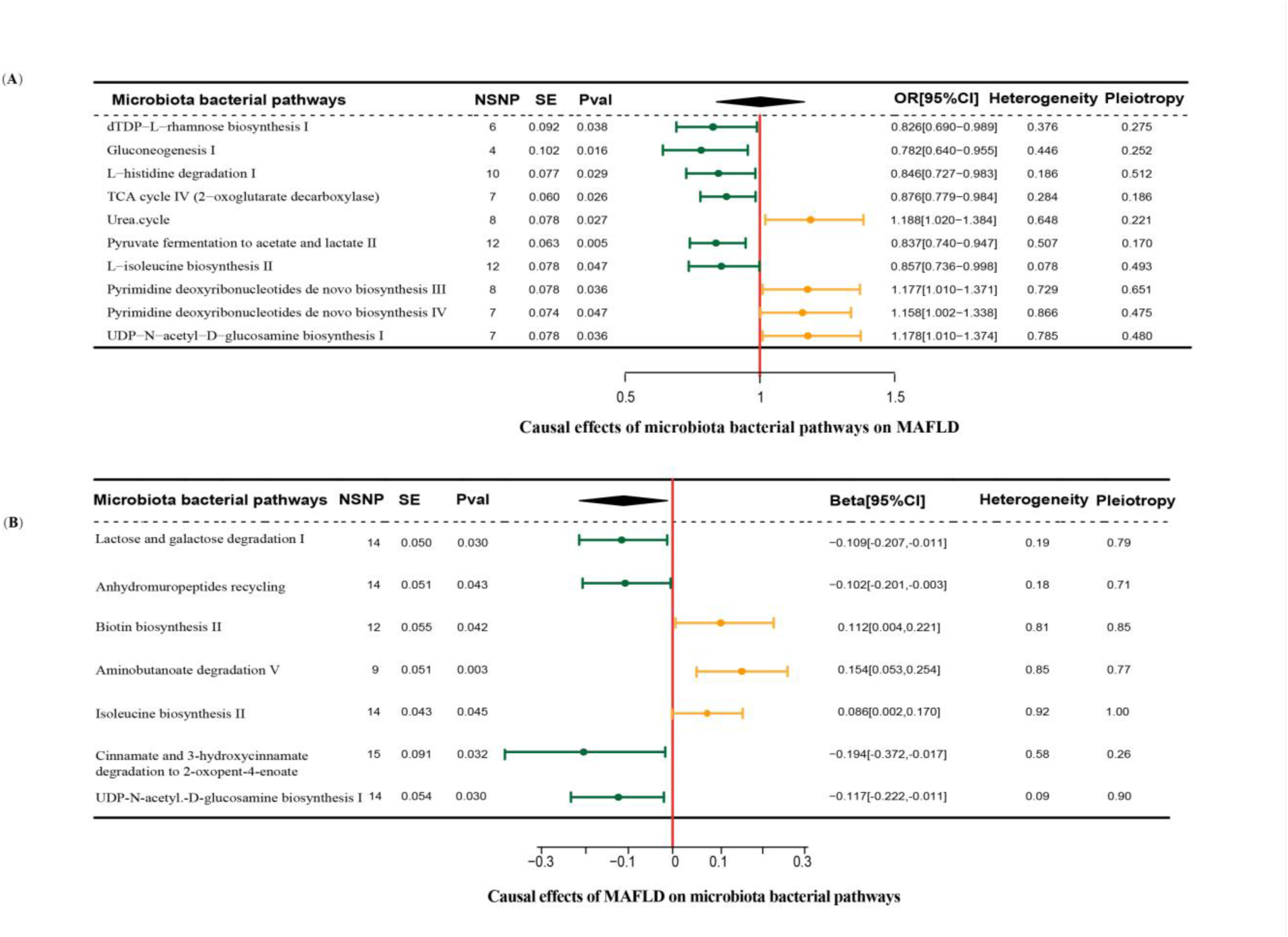
**Forestplot of bidirectional causal effects between bacterial pathways and MAFLD (p < 0.05)**. (A) Causal effects of microbiota bacterial pathways on MAFLD; (B) Causal effects of MAFLD on microbiota bacterial pathways. NSNP, number of SNPs; SE, standard error of coefficient estimate; Pval, p-value of causal estimation in the IVW method; OR [95%CI], odds ratio and 95% confidence interval; Heterogeneity, p-value of heterogeneity analysis; Pleiotropy, p-value of horizontal pleiotropy analysis.

**Supplementary Figure 4.**
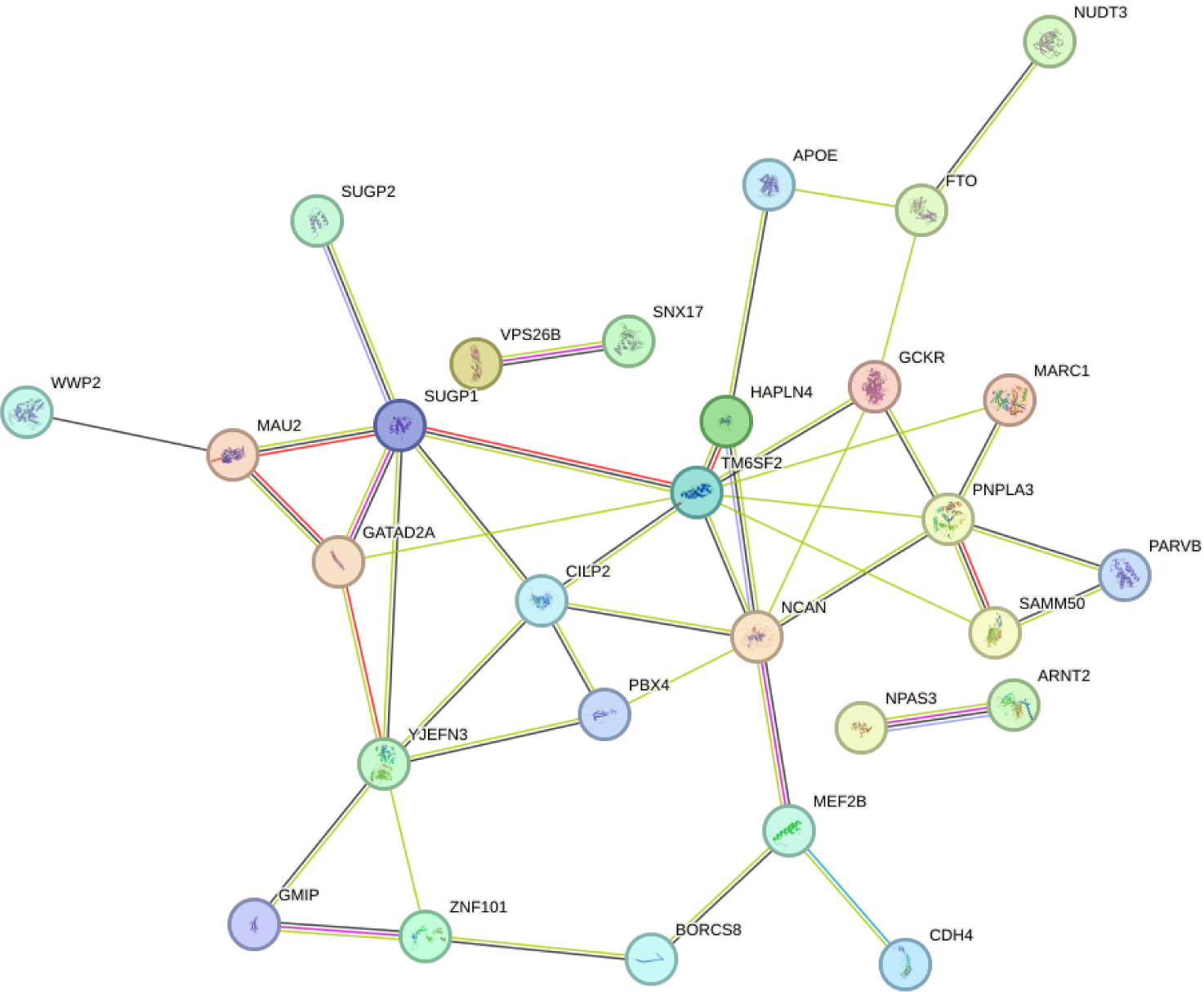
**The Protein-Protein Interaction network analysis of the Lachnospiraceae.bacterium.8.1.57FAA species, the phenylpyruvate, and MAFLD.**

